# Reduced expression of an essential blood-stage *Plasmodium* phosphatidylserine synthase does not modulate parasite resistance to *Pf*ATP4 inhibitors

**DOI:** 10.64898/2026.04.28.721239

**Authors:** Alexis Mann, Montana Sievert, Rubayet Elahi, Shivendra G. Tewari, Krithika Rajaram, Sean T. Prigge

## Abstract

*Plasmodium falciparum* ATP4 mutations A211V and G223R allow parasites to survive the lethal effects of antimalarials PA21A092 (PA92) and cipargamin (CIP), respectively. An A211V mutant line (Dd2^A211V^) treated with PA92 showed enhanced levels of lipid production, which prompted the idea that components of the phospholipid biosynthesis pathway could be involved in the survival mechanism of *Pf*ATP4 mutant parasites. As phosphatidylserine synthase (*Pf*PSS) is the only enzyme that produces phosopholipid phosphatidylserine (PS) in *P. falciparum* parasites, we hypothesized that *Pf*PSS is both essential for parasite survival and that reduced *PfPSS* expression would cause resistant *Pf*ATP4 mutant parasites to become susceptible to PA92 or CIP. We created a CIP-resistant G223R mutant line (Dd2^G223R^) via CRISPR-Cas9 and integrated a conditional *Pf*PSS knockdown construct into a Dd2^A211V^ (↓PSS-Dd2^A211V^) and our Dd2^G223R^ line (↓PSS-Dd2^G223R^). We treated these knockdown lines with PA92 or CIP to determine the half-maximal effective concentration (EC_50_) of each antimalarial with normal or reduced *Pf*PSS levels. While we found that *Pf*PSS is essential for parasite survival, we did not find any significant alterations to the EC_50_ values of PA92 or CIP based on the reduced levels of *Pf*PSS in our mutant lines. Although *Pf*PSS does not appear to be involved, other components of the phospholipid production pathway could still affect the resistance mechanism of *Pf*ATP4 mutations. Identification of novel targets to counteract the mechanism by which *Pf*ATP4 mutant parasites resist lethal drug effects is crucial for the successful application of antimalarials in endemic countries where resistance is on the rise.

## Introduction

Malaria remains a public health concern with nearly 282 million cases and 610,000 associated deaths reported worldwide in 2024 (1). Artemisinin-based combination therapies (ACTs) have been the frontline treatment against these cases in malaria-endemic regions (1). However, the emergence of mutations in the human-infecting parasite *Plasmodium falciparum* that confer partial resistance to ACTs has resulted in cases of ineffective malaria treatment (1). Consequently, there is a pressing need for alternative drug targets and therapies to treat malaria.

One such alternative drug target is the P-type ATPase *Pf*ATP4, identified in chemical drug screens against the chloroquine-sensitive strain *Pf*3D7 and the multidrug-resistant strain *Pf*Dd2 (2-4). *Pf*ATP4 is a Na^+^ efflux/H^+^ influx pump that is crucial for maintaining low cytosolic Na^+^ concentrations in the parasite (5-9). This Na^+^ extrusion activity helps maintain both cellular ionic homeostasis and the integrity of the plasma membrane, which allows for parasite survival (7, 10).

Several classes of antimalarial drugs, including spiroindolones and pyrazoleamides, are thought to inhibit the essential activity of *Pf*ATP4 by binding directly to *Pf*ATP4 or interacting with membrane-associated regulators of *Pf*ATP4 activity (7). Inhibition of *Pf*ATP4 results in an increased concentration of cytosolic Na^+^ in the parasite followed by lethal downstream effects such as cytosolic alkalinization, reduction of cholesterol extrusion and increased membrane rigidity, premature schizogony and eryptosis, and osmotic imbalance and disrupted lipid homeostasis leading to parasite swelling (6-8, 11-20). The accumulation of Na^+^ within the parasite cytosol that results in osmotic imbalance and cell swelling is a unique effect conferred only by *Pf*ATP4-targeting compounds (6, 8, 11, 13, 15, 16, 20, 21).

Parasites exposed to sublethal concentrations of spiroindolones and pyrazoleamides in a lab setting developed drug-resistance mutations (17, 19, 22, 23). For example, the *Pf*ATP4 mutation Ala211Val (A211V) arose following continuous treatment of blood-stage parasites with sublethal levels of a pyrazoleamide, PA21A092 (PA92) while the *Pf*ATP4 mutation Gly223Arg (G223R) appeared following continuous pressure with a spiroindolone, cipargamin (CIP) (18, 23). These mutations localize near the Na^+^ binding cavity of *Pf*ATP4 on the transmembrane domain, which likely reduces the binding affinity of these drugs to *Pf*ATP4 or blocks other inhibitory interactions of these antimalarials with *Pf*ATP4 (5, 6, 10, 17). Of particular interest, the A211V mutation results in both PA92 resistance and CIP hypersensitivity suggesting the existence of important differences in the binding sites of these two drugs (19, 30).

As observed in susceptible parasites treated with *Pf*ATP4-targeting antimalarials, resistant mutants treated with these antimalarials also tend to exhibit slightly impaired Na^+^ regulation (increased intracellular Na^+^ levels) and phospholipid accumulation (9, 17, 18, 19). This phenomenon has been most clearly observed by a genome-scale metabolic network modeling analysis for a PA92-resistant, CIP-hypersensitive Dd2^A211V^ line treated with PA92. Based on time-resolved transcriptomic and metabolomic data, the PA92-treated Dd2^A211V^ line showed an increase in phospholipid synthesis reactions, particularly associated with the production of phosphatidylinositol (PI), phosphatidylcholine (PC), and phosphatidylserine (PS), compared to untreated parental Dd2 parasites (19). However, unlike susceptible parasites, these mutant parasites are somehow able to escape the deleterious effects of disrupted Na^+^ homeostasis (9).

As enhanced phospholipid production is a prominent feature in parasites treated with *Pf*ATP4-targeting antimalarials, we hypothesized that phospholipid biosynthesis participates in the resistance phenotype of these mutants. Blood-stage parasites can synthesize lipids *de novo* from available metabolites or scavenge lipids from exogenous sources like host plasma and red blood cells (RBCs) (24). PS can only be generated through the conversion of exogenous serine via phosphatidylserine synthase (*Pf*PSS) (24). Phosphatidylethanolamine (PE) can be produced *de novo* through either the Kennedy pathway via exogenous ethanolamine or by way of PS decarboxylase (*Pf*PSD) that catalyzes the decarboxylation of PS to form PE (24-28). PC can similarly be produced through the Kennedy pathway with exogenous choline or through the parasite’s phosphoethanolamine methyltransferase (*Pf*PMT) that converts phosphoethanolamine into phosphocholine, an intermediate of the PC biosynthetic pathway (24, 26). As *Pf*PSS is the only enzyme capable of producing PS, which can then be utilized to generate other lipids for the cell, we pinpointed this enzyme as a target of interest for our antimalarial assays. Our choice was further supported by a study in related apicomplexan parasite *Toxoplasma gondii* in which the loss of *PSS* expression resulted in parasite death, indicating that PSS activity is essential for parasite survival (29).

Consequently, we aimed to determine if the increased levels of PS production observed in Dd2^A211V^ parasites treated with the *Pf*ATP4 inhibitor PA92 play a role in drug resistance. We hypothesized that reducing PS production would increase parasite susceptibility to PA92. In contrast, the CIP-hypersensitive Dd2^A211V^ parasites did not trigger increased PS production with CIP treatment (19). We further hypothesized that PS production could instead play a role in CIP-resistant parasites. To this end, we generated CIP-resistant *Pf*ATP4 G223R mutant line (Dd2^G223R^). Then, to determine whether reducing PS production would increase parasite susceptibility to a *Pf*ATP4-inhibiting antimalarial, we knocked down *Pf*PSS expression in mutant lines Dd2^A211V^ and Dd2^G223R^ and treated the parasites with PA92 or CIP respectively. Although we were able to demonstrate that *Pf*PSS activity is essential for parasite survival in our mutant lines, we were unable to identify a significant shift in drug susceptibility, determined by half-maximal effective concentration (EC_50_), to the *Pf*ATP4-targeting antimalarials PA92 or CIP in parasites with reduced *Pf*PSS levels.

Further investigation into *Pf*PSS and its potential connection to *Pf*ATP4-targeting antimalarials is needed, but our initial findings emphasize the importance of phospholipid biosynthesis for parasite survival that could be exploited for novel drug targets. Additionally, other enzymes associated with phospholipid production should be studied to fully understand the resistance mechanism of emerging *Pf*ATP4 mutations observed in endemic countries.

## Results

### *Generation of the CIP-resistant line* Dd2^G223R^

Results from a genome-scale metabolic network modeling analysis integrating time-resolved transcriptomic and metabolomic data showed that PA92-treated Dd2^A211V^ parasites exhibited upregulated lipid metabolism, including enhanced biosynthesis of phospholipids such as PS, compared to untreated parasites (19). To study this response to drug treatment, we aimed to investigate whether ablating PS synthesis by knockdown of *Pf*PSS levels in two blood-stage, *Pf*ATP4 mutant parasite lines, Dd2^A211V^ and Dd2^G223R^, would increase the susceptibility of the mutant parasites to *Pf*ATP4-targeting antimalarials.

The Dd2^A211V^ isogenic mutant line was previously generated and characterized for a resistance phenotype against PA92 by the Vaidya lab (19, 30). However, a Dd2^G223R^ isogenic mutant line that conveys resistance to CIP was not available. We therefore used markerless genome editing to generate a Dd2^G223R^ parasite line (**Figure 1A, Supplemental Table 1**). We validated the resulting Dd2^G223R^ line with Sanger sequencing (**Figure 1B, Supplemental Figure 1A**). Then, we evaluated the resistance phenotype of Dd2^G223R^ parasites by determining the EC_50_ of CIP. We observed a ∼14-fold increase of the CIP EC_50_ value in Dd2^G223R^ parasites compared to the isogenic Dd2 parental line (p = 0.0175) (**Figure 1C, Supplemental Figure 1B**). For comparison, a ∼6-fold increase in EC_50_ was reported for a G223R mutant line that was generated via continuous spiroindolone drug pressure (23).

**Figure 1.**
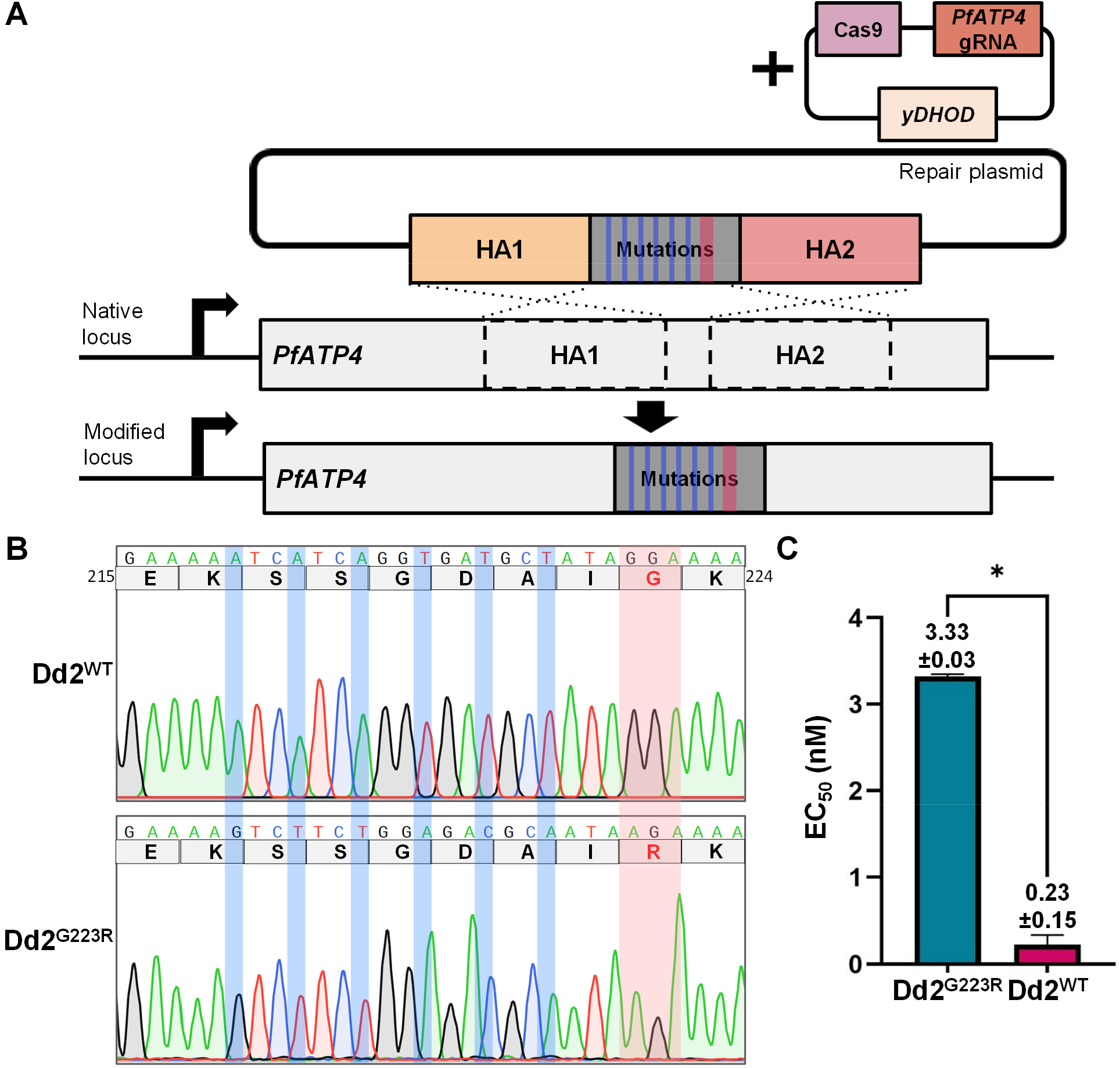
Generation of the CIP-resistant line Dd2^G223R^. A) Diagram depicting the incorporation of the G223R mutation into the *PfATP4* locus. Briefly, a Cas9 endonuclease directed by a guide RNA (gRNA) cleaves the native *PfATP4* locus. The resulting double-stranded break is repaired by a double crossover homologous recombination event directed by the two homology arms (HAs) of the repair plasmid. The HAs flank a segment of mutated DNA that includes a glycine to arginine (G → R) mutation (light red) and shield mutations (light blue). The modified *PfATP4* locus then translates to a *Pf*ATP4 protein containing a single amino acid replacement (G223R). *yDHOD*, yeast dihydroorotate dehydrogenase. B) Chromatogram of the DNA and amino acid sequence of the Dd2^G223R^ mutant line compared to the native (Dd2^WT^) sequence. Each codon preceding the desired glycine to arginine (G → R) amino acid replacement (light red) in Dd2^G223R^ contained a synonymous shield mutation (light blue) to prevent Cas9 cleavage of the mutated *PfATP4* sequence while maintaining the same amino acid sequence as the Dd2^WT^ line. C) CIP EC_50_ values for Dd2^WT^ and Dd2^G223R^ mutant parasites. The EC_50_ values were significantly different (p=0.0175) with the mutant line presenting an EC_50_ approximately 14-times greater than the control line. Mean EC_50_ ± standard deviation values are shown. Error bars represent the standard error of the mean for two biological replicates, each conducted in quadruplicate. Statistical significance was determined using a two-tailed unpaired t-test with Welch’s correction.

### *Generation of* ↓PSS-Dd2^A211V^ and ↓PSS-Dd2^G223R^ *parasites*

*Pf*PSS is the only PS synthase present in *P. falciparum* and, consequently, plays an important role in the appropriate maintenance of phospholipid homeostasis in the parasite (25). As with its mammalian ortholog PSS2, *Pf*PSS catalyzes a base exchange reaction, replacing the ethanolamine group of PE with an exogenously supplied serine group to generate PS (25, 31). To ablate PS production, we used CRISPR-Cas9 mediated gene editing to modify the 3’ untranslated region (UTR) of native *PfPSS* in both the Dd2^A211V^ and Dd2^G223R^ background. Of note, the modified *PfPSS* locus encodes a C-terminal 3× FLAG tag, a 10× aptamer array, and the TetR-DOZI fusion protein (**Figure 2A, Supplemental Table 1**). The FLAG tag allows for protein visualization, and the TetR-DOZI aptamer machinery allows for inducible control over protein translation (32). Briefly, the presence of the small molecular anhydrotetracycline (aTc) in the parasite media promotes protein production (+PSS) while the removal of aTc from the media results in the knockdown of protein translation (-PSS). The parasite lines that express the *Pf*PSS knockdown system will hereafter be referred to as ↓PSS-Dd2^A211V^ and ↓PSS-Dd2^G223R^. We validated the successful generation of ↓PSS-Dd2^A211V^ and ↓PSS-Dd2^G223R^ via genotyping PCR (**Figure 2B-C, Supplemental Table 1**). We additionally submitted ↓PSS-Dd2^G223R^ for whole genome sequencing (WGS) to verify the presence of the *Pf*ATP4 G223R mutation in this knockdown line.

**Figure 2.**
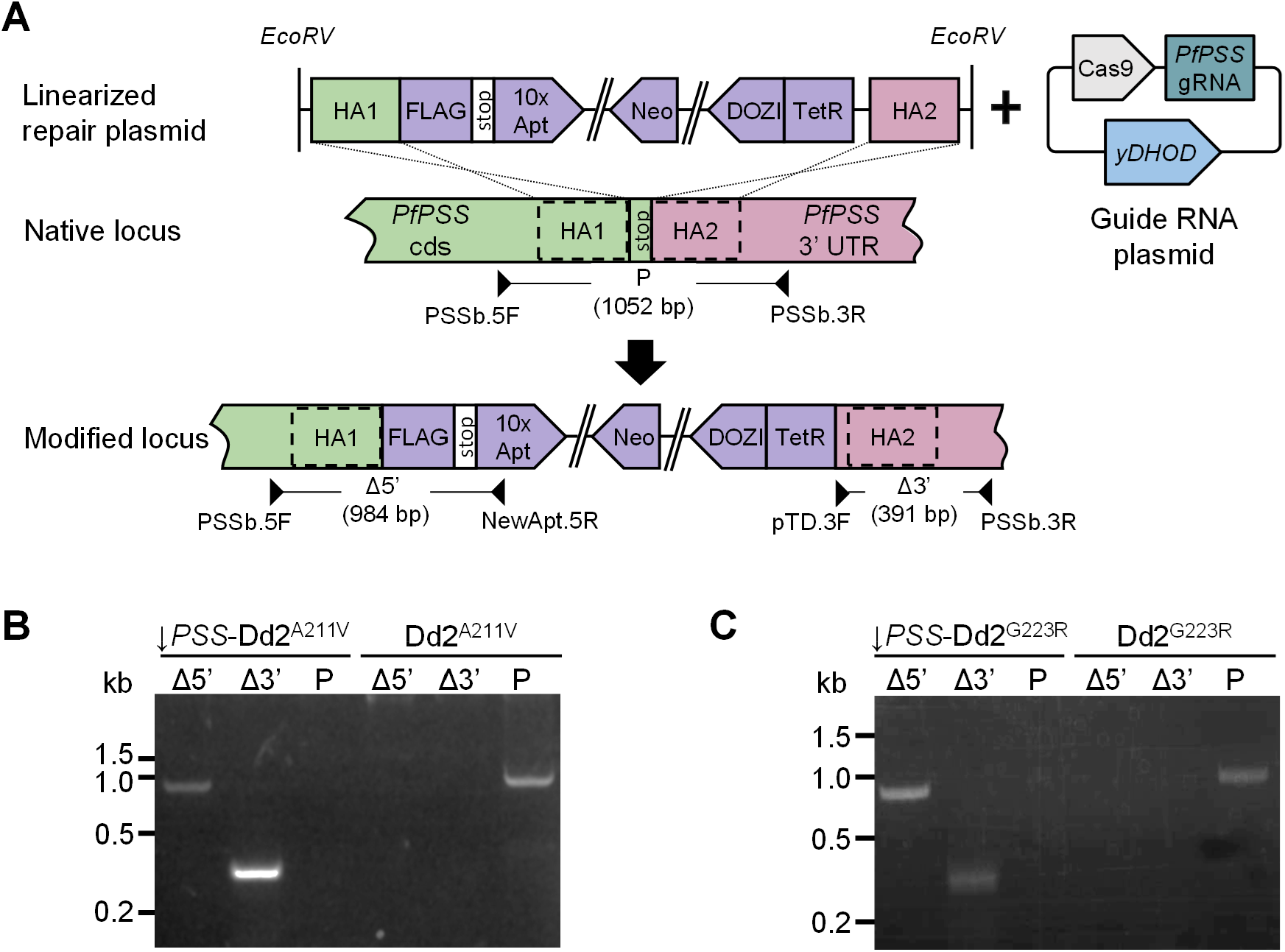
Generation of ↓PSS-Dd2^A211V^ and ↓PSS-Dd2^G223R^ parasites. A) Diagram depicting the incorporation of *Pf*PSS knockdown machinery into the native *PfPSS* locus through a double crossover homologous recombination event. Briefly, a guide RNA (gRNA) directs a Cas9 endonuclease to introduce a double-stranded break in the native *PfPSS* locus. This cleavage is repaired by homologous recombination directed by the two homology arms (HAs) of the *Eco*RV-linearized repair plasmid. The plasmid includes FLAG epitope tags for visualization of the *Pf*PSS protein and the TetR-DOZI/aptamer system for conditional *Pf*PSS knockdown. The location and direction of the primers used for integration confirmation of the knockdown machinery are shown as black arrowheads. Solid black lines between the primer pairs show the expected sizes of the genotyping PCR products. B) Genotype confirmation gel showing the successful integration of the *Pf*PSS knockdown machinery into the ↓PSS-Dd2^A211V^ or C) ↓PSS-Dd2^G223R^ line compared to the parental Dd2^A211V^ or Dd2^G223R^ lines, respectively.

### *Pf*PSS *is essential for parasite growth and survival*

A genome-wide *piggyBac* mutagenesis study in *P. falciparum* predicted *PfPSS* to be essential for parasite survival; however, targeted reverse genetic studies of *PfPSS* have not been performed (33, 34). To test the essentiality of *PfPSS*, we cultured our ↓PSS-Dd2^A211V^ and ↓PSS-Dd2^G223R^ lines either with or without aTc over the course of eight days (**Figure 3A**). For both mutant lines, parasites cultured without aTc displayed significantly reduced growth by Day 4, with parasites undetectable by flow cytometry by Day 6 (**Figure 3B-C, Supplemental Figure 2A-B**). For the first four days of the growth assay, we also collected samples for complementary Western blots to validate protein knockdown (**Figure 3A**). The Western blots validated the loss of *Pf*PSS protein production in both mutant lines as the presence of FLAG-tagged PSS was lost within two days of removing aTc from the parasite culture (**Figure 3D-E, Supplemental Figure 2C-D**). Although the expected size of *Pf*PSS is 42 kDa, we consistently observed protein bands at 32 kDa on our Western blots that were recognized by anti-FLAG antibodies and were responsive to aTc treatment (31). This phenomenon could be due to amino-terminal processing of the protein, resulting in a smaller protein that still retains the carboxy-terminal FLAG tag. Alternatively, since we do not observe any unprocessed protein, intact *Pf*PSS could migrate at an anomalous rate on the gel. In either case, the loss of *Pf*PSS was quickly followed by significantly reduced growth after the second growth cycle in both our ↓PSS-Dd2^A211V^ and ↓PSS-Dd2^G223R^ lines. Thus, *Pf*PSS protein is essential for parasite growth and survival in both of our *Pf*ATP4-mutated lines.

**Figure 3.**
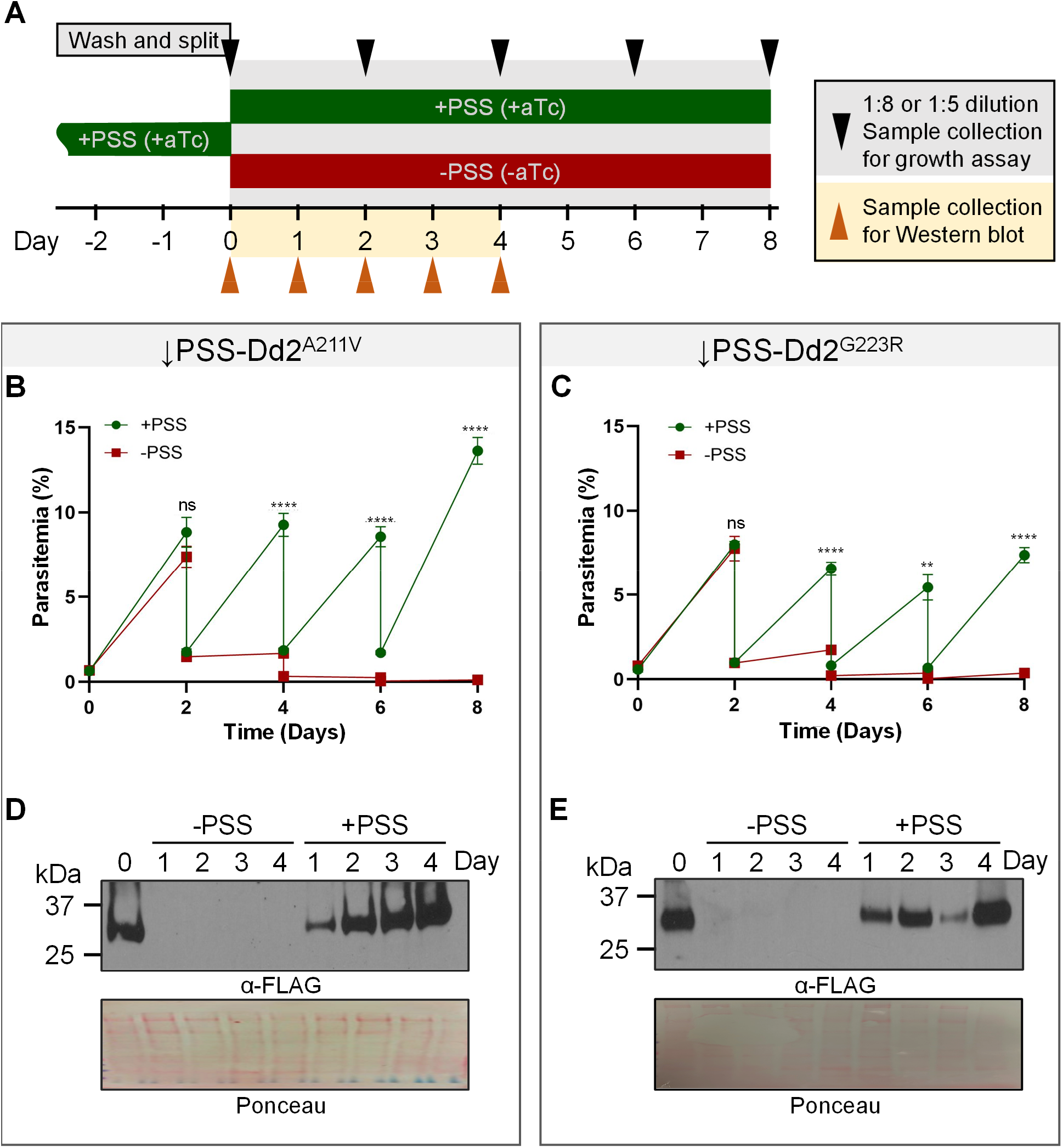
*Pf*PSS is essential for parasite growth and survival. A) Timeline of sample collection for growth assay and Western blot. Both ↓PSS-Dd2^A211V^ and ↓PSS-Dd2^G223R^ were kept on anhydrotetracycline (aTc) to maintain *Pf*PSS protein levels. On Day 0 of the experiment, parasites were washed of aTc and split equally to be cultured either with aTc (normal PSS levels, +PSS) or without aTc (reduced PSS levels, -PSS). Each line in both conditions was cultured for eight days. Every day from Days 0-4, samples were collected for Western blot. Samples were also collected every other day for flow cytometry to monitor growth and were cut back either 1:5 for ↓PSS-Dd2^A211V^ or 1:8 for ↓PSS-Dd2^G223R^ to prevent overgrowth. B) Graph depicting parasite growth for ↓PSS-Dd2^A211V^ or C) ↓PSS-Dd2^G223R^ cultured with aTc (+PSS) or without aTc (-PSS) over the course of eight days. Parasites cultured with aTc have endogenous *Pf*PSS levels and grow with consistent culture cut-backs every two days. Parasites cultured without aTc have reduced *Pf*PSS levels and, by day four without aTc, begin showing statistically significant growth defects that result in undetectable parasite growth compared to those parasites grown in the presence of aTc. Error bars represent the standard error of the mean of two biological replicates, each conducted in quadruplicate. Statistical significance was determined using a two-way ANOVA with Šídák’s multiple comparisons test. ns: not significant (p>0.05), ** significant (p=0.0021), **** significant (p<0.0001). D) Western blot of the ↓PSS-Dd2^A211V^ or E) ↓PSS-Dd2^G223R^ line cultured either with aTc (+PSS) or without aTc (-PSS) over the course of four days. Bands for PSS, approximating 30 kDa, are visible when aTc is present (+PSS), but disappear by Day 1 following the removal of aTc from the parasite culture (-PSS). The FLAG-tagged PSS bands were identified following incubation with an α-FLAG primary antibody and the loading control was determined via whole-protein Ponceau staining.

### *Reduced Pf*PSS *production does not alter the susceptibility of* ↓PSS-Dd2^A211V^ *and* ↓PSS-Dd2^G223R^ *parasites to PA92 and CIP treatment*

Finally, we wanted to determine whether the reduction of *Pf*PSS protein levels affected the ability of our *Pf*ATP4 mutant parasite lines to grow in the presence of the antimalarial drugs PA92 or CIP. We predicted that *Pf*PSS knockdown in the ↓PSS-Dd2^A211V^ and ↓PSS-Dd2^G223R^ lines would result in enhanced drug sensitivity (a lower EC_50_) similar to what was observed in the Dd2 parental parasites. To test this prediction, two days after initiating *Pf*PSS knockdown by aTc washout, we treated ↓PSS-Dd2^A211V^ with a dilution series of PA92 and ↓PSS-Dd2^G223R^ with a dilution series of CIP for three days to determine their respective EC_50_ values (**Figure 4A**). We did not find a statistically significant difference between the EC_50_ values of either mutant line between normal (+PSS) and reduced *Pf*PSS (-PSS) levels (**Figure 4B-C, Supplemental Figure 3A-B)**. Consequently, we concluded that the loss of *Pf*PSS and, subsequently, PS does not alter the susceptibility of ↓PSS-Dd2^A211V^ to PA92 or ↓PSS-Dd2^G223R^ to CIP.

**Figure 4.**
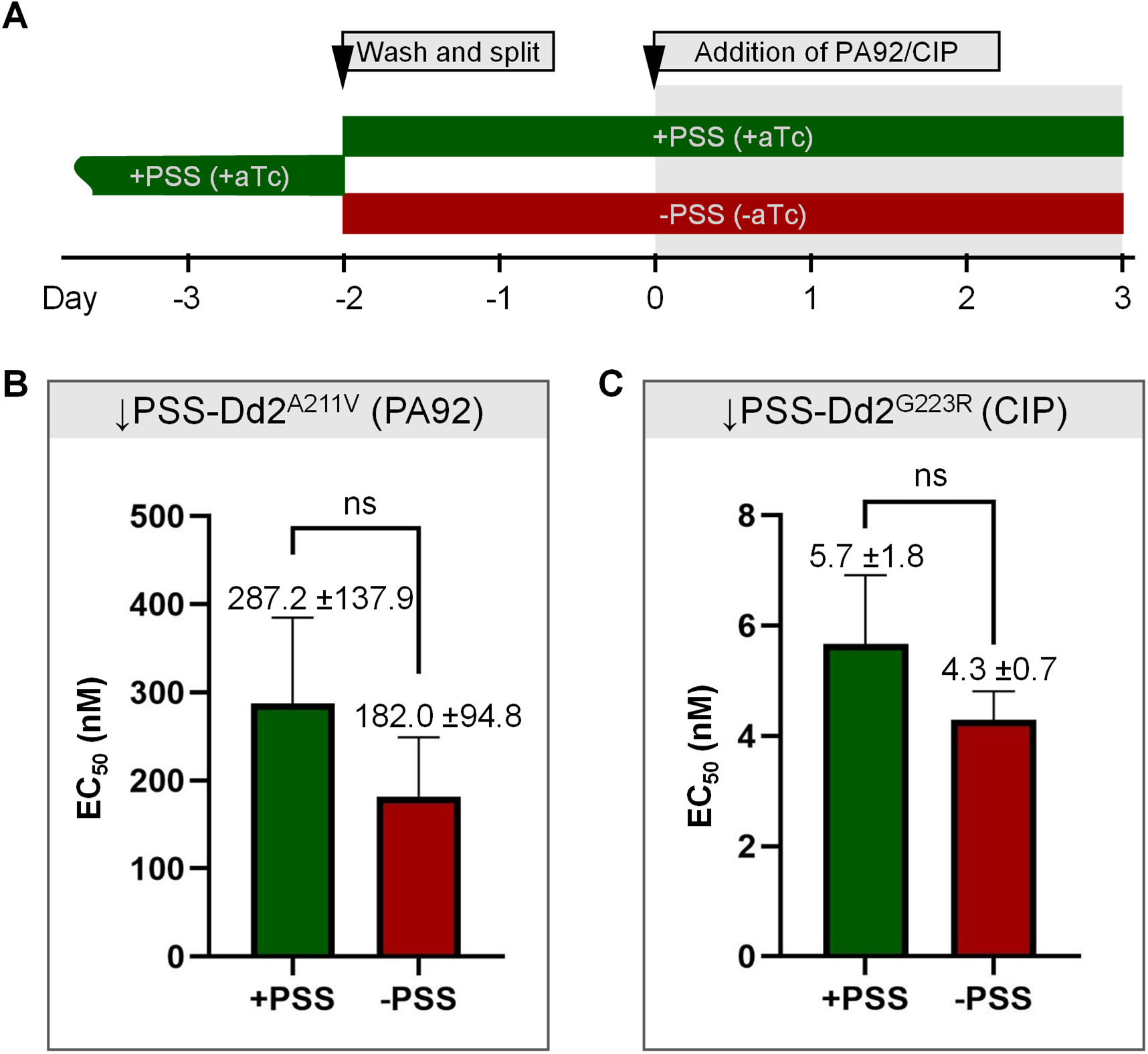
Reduced *Pf*PSS production does not alter the susceptibility of ↓PSS-Dd2^A211V^ and ↓PSS-Dd2^G223R^ parasites to PA92 and CIP treatment. A) Schematic of EC_50_ assay setup. Both ↓PSS-Dd2^A211V^ and ↓PSS-Dd2^G223R^ were kept on aTc to maintain *Pf*PSS levels. Two days before the experiment began, the cultures were washed of aTc and split equally to be cultured either with aTc (normal PSS levels, +PSS) or without aTc (reduced PSS levels, -PSS). On Day 0 of the experiment, the ↓PSS-Dd2^A211V^ line cultured in both conditions was introduced to a dilution series of PA92 and ↓PSS-Dd2^G223R^ cultured in both conditions was introduced to a dilution series of CIP. Samples were then collected from all conditions after three days on their respective drug. B) EC_50_s of ↓PSS-Dd2^A211V^ cultured either with normal or reduced PSS levels in the presence of a PA92 dilution series, or C) ↓PSS-Dd2^G223R^ cultured either with normal or reduced PSS levels in the presence of a CIP dilution series. There was no significant difference (p>0.05) between the EC_50_s of parasites cultured without aTc (reduced PSS) and those cultured with aTc (normal PSS) when in the presence of *Pf*ATP4 inhibiting antimalarials. Mean EC_50_ values ± the standard deviation are shown for the experimental and parental groups. Error bars represent the standard error of the mean from two biological replicates, each conducted in quadruplicate. Statistical significance was determined using a two-tailed unpaired t-test with Welch’s correction.

## Discussion

In this study, we used Dd2^A211V^ and Dd2^G223R^, two mutant lines resistant to *Pf*ATP4 inhibitors, to study i) the essentiality of *PfPSS* for parasite survival and ii) whether *Pf*PSS plays a role in the resistance mechanism associated with *Pf*ATP4 mutations. In other organisms, PSS catalyzes the production of PS and is associated with maintaining the integrity of both plasma and organellar membranes (35, 36). Similarly, malaria parasites must generate high amounts of phospholipids, 4-8% of the total pool consisting of PS, to form the plasma membranes and internal membranous compartments of the many daughter cells produced during schizogony (37, 38). PSS from the related apicomplexan parasite *T. gondii* likely shares this function because the loss of *Tg*PSS resulted in the disruption of membrane integrity and parasite death (29). Based on the critical role PS plays in membrane maintenance and the essentiality of *PSS* in other apicomplexan organisms, we hypothesized that the reduction of *Pf*PSS protein levels would result in parasite death – a phenomenon we did observe in our *Pf*ATP4 mutant lines. Consequently, we concluded that *Pf*PSS and its role in PS production are essential for the survival of our *Pf*ATP4 mutant *Plasmodium* parasites.

We then interrogated whether *Pf*PSS could contribute to the resistance phenotype that our *Pf*ATP4 mutant lines display toward *Pf*ATP4-targeting antimalarials. *Plasmodium* parasites treated with *Pf*ATP4-targeting antimalarials consistently experience disrupted ionic homeostasis across the plasma membrane, which can lead to enhanced phospholipid synthesis, cell swelling, and an increased number of daughter cells produced during replication (15, 18, 19). These downstream effects of antimalarial treatment have specifically been observed in the PA92-resistant, CIP-hypersensitive *Pf*ATP4 mutant Dd2^A211V^; however, this resistant line does not undergo lysis and parasite death, hallmarks of the typical treatment of susceptible parasites (8, 19, 20). As enhanced lipid biosynthesis is a common feature of treatment with *Pf*ATP4-targeting antimalarials, we hypothesized that *Pf*PSS, an enzyme associated with phospholipid production, might help *Pf*ATP4 mutant parasites evade the deleterious downstream effects of *Pf*ATP4-targeting antimalarials. Contrarily, we did not detect significantly different inhibitory capacities of *Pf*ATP4-targeting antimalarials used to treat resistant lines with different levels of *Pf*PSS. From this preliminary analysis, *Pf*PSS does not appear to play a role in mediating resistance of *Pf*ATP4 mutant parasites to their respective *Pf*ATP4-targeting antimalarial.

Since *PfPSS* is essential, our antimalarial susceptibility experiments had to be carried out in gene knockdown rather than knockout lines to ensure we were i) able to create a viable line and ii) have at least a short window of time to study the effects of reduced *Pf*PSS protein levels. We began drug treatment on Day 2 after initiating *Pf*PSS knockdown because the parasites were still growing even as *Pf*PSS levels were largely reduced. The parasites could have still produced some amount of *Pf*PSS, albeit at low levels that were undetectable in our Western blot assays. As the reduced level of *Pf*PSS protein achieved by our knockdown method did not support long-term parasite growth, it appears equally unlikely such a diminished protein pool could mediate the resistance mechanism of *Pf*ATP4 mutations against PA92 or CIP treatment.

Although *Pf*PSS and PS do not appear to be involved in the resistance mechanism conferred by *Pf*ATP4 mutations, other components of the lipid biosynthesis pathway have also exhibited altered production in response to drug treatment and merit further study. Susceptible parasites treated with PA92 specifically showed enhanced production of myoinositol phosphate phosphatase (MIPP) and myoinositol 3-phosphate synthase (MIPS), both of which are directly involved in the production of myoinositol, a precursor for phosphatidylinositol (PI) (18). Myoinositol has been long associated with osmoregulatory functions in various organisms (39-41). PI itself is a structural lipid associated with membrane trafficking while its phosphorylated counterpart, phosphatidylinositol 3-phosphate (PI3P), is largely involved with lipid trafficking (19, 42). These lipids and lipid-associated enzymes could serve to redirect the extensive phospholipid pool produced in response to *Pf*ATP4-targeting antimalarial treatment toward parasite proliferation rather than death in mutant lines. Consequently, future experiments could focus their efforts on myoinositol- or PI-producing enzymes to determine if these components of lipid biosynthesis play a role in mediating the resistance conferred by *Pf*ATP4 mutant parasites against *Pf*ATP4-targeting antimalarials.

Overall, we determined that *PfPSS* is essential for the survival of *Pf*ATP4 mutant parasites but does not appear to be involved in modulating resistance conferred by *Pf*ATP4 mutations. Future work should instead target other lipid-producing enzymes with a focus on investigating the lipid landscape that arises following *Pf*ATP4 inhibitor treatment and loss of target gene expression. Such studies will help guide the use of *Pf*ATP4 inhibitors in combination with available antimalarial therapies to minimize the threat of emerging drug resistance.

## Materials and Methods

### Asexual blood-stage culture and maintenance

All experiments were performed in a *P. falciparum* Dd2 strain. Asexual blood-stage *P. falciparum* parasites were cultured in human O^+^ RBCs at 1% hematocrit in complete medium with AlbuMAX II (CMA). CMA was made from RPMI 1640 medium with L-glutamine (Catalog #R8999, USBiological Life Sciences) supplemented with 20 mM HEPES (Catalog #H4034, MilleporeSigma), 0.2% sodium bicarbonate (Ref #25080-094, ThermoFisher Scientific), 12.5 μg/mL hypoxanthine (Catalog #H9377, MilleporeSigma), 5 g/L Albumax II (Ref #11021-045, ThermoFisher Scientific), and 25 μg/mL gentamicin (Catalog #G3632, MilleporeSigma). Cultures were maintained in 25 cm^2^ gassed flasks (94% N_2_, 3% O_2_, 3% CO_2_) and incubated at 37°C.

### Construction of transfection plasmids

To generate the *Pf*ATP4 (PF3D7_1211900) G223R mutation in Dd2 parasites, a repair plasmid (pRSng-G223R) and a corresponding guide RNA (gRNA) plasmid (pCasG-G223R) were constructed (32, 43). The pRSng-G223R repair plasmid was generated from a synthetic DNA fragment cloned into a pUC57 backbone (LifeSct). Flanked by NotI sites, the synthetic DNA contained two ∼350 bp *PfATP4* homology arms (HAs) positioned upstream and downstream of a modified *PfATP4* coding sequence. The wild-type *PfATP4* nucleotide sequence (‘aaa tca tca ggt gat gct ata gga’) was modified (‘aag tct tct gga gac gca ata aga’) to introduce a ‘gga’ → ‘aga’ single nucleotide polymorphism (SNP) resulting in the G223R amino acid substitution.

Additionally, synonymous shield mutations were introduced into the codons upstream of the G223R codon to prevent Cas9 cleavage of the mutated sequence while preserving the native amino acid sequence (44). The synthetic fragment was excised from pUC57 by digestion with *Not*I (Catalog #R3189, New England Biolabs) and ligated into the NotI site of the pRSng backbone with T4 DNA ligase (Catalog #M0202, New England Biolabs) to form the repair plasmid pRSng-G223R (32).

The 20 bp gRNA sequence (‘aaa tca tca ggt gat gct at’) targeting the *PfATP4* G223 position was synthesized as complementary forward and reverse oligonucleotides flanked by 17 bp sequences homologous to the pCasG plasmid insertion site (**Supplemental Table 1**). The oligos were annealed and cloned into the BsaI sites of the pCasG-LacZ plasmid using ligase independent cloning (In-Fusion) (Catalog #639650, Clontech Laboratories) to generate the pCasG-G223R construct.

To generate *Pf*PSS (PF3D7_1366800) knockdown (↓PSS) constructs, we used repair plasmid pTDNF that contains a TetR-DOZI expression cassette (45). First, HAmcs, a 314 bp long adapter flanked by AscI and AatII restriction enzyme recognition sites, was synthesized (Twist Biosciences). This 314 bp long adapter includes an EcoRV, a XhoI, and 2 BsaI restriction enzyme recognition sites to facilitate cloning of the HAs and linearization of the pTDNF-↓PSS plasmid prior to transfection. The adapter was also recodonized to introduce shield mutations in the gRNA binding site to prevent Cas9-mediated cleavage in the transgenic parasites. Then, the synthetic HAmcs was digested by *Asc*I (Catalog #R0558, New England Biolabs) and *Aat*II (Catalog #R0117, New England Biolabs) for ligation into the equivalent restriction sites of pTDNF-PfDHO to generate the pTDNF-HAmcs plasmid (45). Finally, HA1 and HA2 were amplified from parental parasite lysates (**Supplemental Table 1**). The HA1 PCR amplicon was ligated into the BsaI sites of pTDNF-HAmcs to generate pTDNF-PSS.HA1, and the HA2 amplicon was digested by *Asc*I and *Bam*HI (Catalog #R3136, New England Biolabs) preceding insertion into the relevant sites of pTDNF-PSS.HA1 to generate the pTDNF-↓PSS plasmid.

The 20 bp gRNA sequence (‘ctt acc aga aat aaa acc ta’) targeting *PfPSS* was synthesized as complementary 5’-phosphorylated forward and reverse oligos (**Supplemental Table 1**). The oligos were annealed and cloned into the BsaI sites of pCasG-LacZ plasmid using T4 DNA ligase to generate the pCasG-↓PSS plasmid (43).

### Parasite transfections

To generate the Dd2^G223R^ line, 400 µL of uninfected human RBCs were electroporated using a Bio-Rad Gene Pulser Xcell system (0.31 kV, 925 µF) with 65 µg each of the pRSng-G223R plasmid and the pCasG-G223R plasmid. Following electroporation, Dd2-infected RBCs were mixed with the plasmid-containing RBCs and cultured in CMA for 48 hrs. After 48 hrs, the transfected parasites were cultured with 1.2 nM CIP (a gift from Dr. Akhil Vaidya) for 6 days to select for the G223R *PfATP4* allele before being allowed to recover without drug. CIP-resistant parasites were detectable via thin smear Giemsa staining approximately 18 days post-transfection and were further validated with Sanger sequencing for the presence of the G223R mutation. Furthermore, to ensure the Dd2^G223R^ line was free of WT G223 contamination, the parasites were cloned via limiting dilution and re-sequenced to be used in subsequent experiments.

To generate ↓PSS lines, the ↓PSS-pTDNF repair plasmid was linearized with *Eco*RV (Catalog #R3195, New England Biolabs). Linearized ↓PSS-pTDNF and the pCasG-↓PSS plasmid were co-transfected into RBCs as described above. Following transfection, these RBCs were introduced to either the Dd2^A211V^ or Dd2^G223R^ parasites and were cultured with CMA supplemented 0.5 µM anhydrotetracycline (aTc) (Item #10009542, Cayman Chemical) for 48 hrs. After 48 hrs, the parasites were continuously cultured with 0.5 mg/mL G418 sulfate (Catalog #11811031, ThermoFisher Scientific) and aTc to select for parasites with successful incorporation of the knockdown construct into the native *PfPSS* locus. After approximately 20 days, transgenic ↓PSS*-*Dd2^A211V^ and ↓PSS*-*Dd2^G223R^ parasites appeared in culture and were validated by genotyping PCRs. Clonal parasites were obtained by limiting dilution in 96-well plates for the ↓PSS*-*Dd2^A211V^ and ↓PSS*-*Dd2^G223R^ parasites. A representative clone for each transgenic line was chosen for all the experiments.

### Genotyping PCR

Lysates from parental (Dd2^A211V^ and Dd2^G223R^) and transgenic (↓PSS*-*Dd2^A211V^ and ↓PSS*-* Dd2^G223R^) parasite cultures were prepared by incubating aliquots of cultures at 90 °C for 5 min. The resulting lysates were used for genotype confirmation. To confirm the insertion of the 3× FLAG tag and the knockdown machinery in the transgenic lines, the 5’- and 3’-ends of the modified *PSS* locus (Δ5’ and Δ3’, respectively) and the parental gene locus (P) were amplified from the transgenic and appropriate parental lines with corresponding primers (**Supplemental Table 1**).

All the cultures used in this study were regularly tested for mycoplasma contamination with an in-house developed diagnostic PCR (**Supplemental Table 1**).

### Growth Assay

To assess whether reduced *Pf*PSS levels affected parasite growth, ↓PSS-Dd2^A211V^ and ↓PSS-Dd2^G223R^ parasites were washed three times to remove any residual aTc. Flat-bottom 96-well plates (Catalog #267427, ThermoFisher Scientific) were then seeded with 200 μL culture/well at 0.5% parasitemia and 1% hematocrit in either the presence or absence of 0.5 µM aTc. Every 48 hrs over a period of 8 days, parasite samples were collected and growth was monitored following SYBR green I (Catalog #S7563, ThermoFisher Scientific) staining using an Attune Nxt Flow Cytometer (ThermoFisher Scientific) as previously described (43). Immediately following sample collection, the cultures were diluted either 1:5 for ↓PSS-Dd2^A211V^ or 1:8 for ↓PSS-Dd2^G223R^ to prevent parasite over-growth. Growth data were collected from two independent biological replicates (each in quadruplicate) and analyzed with Prism V10.6 (GraphPad Software). Statistical significance was determined using a two-way ANOVA with Šídák’s multiple comparisons test.

### Western Blot

Pellets of ↓PSS*-*Dd2^A211V^ and ↓PSS*-*Dd2^G223R^ parasites were harvested by centrifugation at 500 ×g for 5 min. The pellets were treated with 0.03% saponin (Catalog #S4521, MilliporeSigma) in 1× PBS at room temperature to release the parasites from the RBCs. The released parasites were then pelleted by centrifugation at 1940 ×g for 10 min at 4 °C. The pellet was washed twice with 1 mL 1× PBS. The saponin-isolated parasite pellets were then resuspended in 1× lithium dodecyl sulfate (LDS) sample buffer (Catalog #NP007, ThermoFisher Scientific) supplemented with 2% β-mercaptoethanol (BME) (Catalog #M7154, MilleporeSigma) and boiled at 95 °C for 5 min to ensure complete protein denaturation and solubilization.

The parasite lysate was then resolved by sodium dodecyl sulfate polyacrylamide gel electrophoresis (SDS-PAGE) on 4-12% gradient reducing gels (Ref #NPO335BOX, ThermoFisher Scientific) and transferred to a nitrocellulose membrane. Membranes were briefly stained with Ponceau S stain (0.1% in 5% acetic acid) before blocking with 5% non-fat dry milk in 1× PBS containing 0.1% Tween-20 (Milk/PBST) for one hour at room temperature. The membranes were incubated with 1:15,000 mouse anti-FLAG M2 (Catalog #F3165-1MG, MilleporeSigma) primary antibody diluted in milk/1× PBS containing 0.1% Tween-20 (PBST) overnight at 4 °C. The membrane was then washed three times in PBST and incubated with sheep anti-mouse horseradish peroxidase (HRP)-conjugated secondary antibody (Catalog #GENA931, MilliporeSigma) diluted 1:10,000 in milk/PBST for 1 hr at room temperature. The resulting chemiluminescent signals were observed using SuperSignal West Pico chemiluminescent substrate (Catalog #34577, ThermoFisher Scientific) and visualized on autoradiography film (Catalog #BL1-810-100, Stellar Scientific).

### Determination of half-maximal effective concentration (EC_50_)

To determine the EC_50_ of CIP for Dd2 and Dd2^G223R^ parasites, asynchronized parasites were seeded at 1% parasitemia and 1.5% hematocrit in a 96-well flat-bottom plate in quadruplicate. CIP was added to parasites from 1,000× stock solutions dissolved in dimethyl sulfoxide (DMSO) to generate a 3-fold concentration series of CIP (16.9 pM to 1 μM). After 72 hrs incubated at 37 °C in a modular incubator chamber gassed with 3% O_2_, 3% CO_2_, and 94% N_2_, parasite growth was determined following SYBR green I staining via flow cytometry as previously described (43).

To determine whether reduced levels of *Pf*PSS alters the ability of two *Pf*ATP4 mutations (A211V and G223R) to confer resistance to PA92 and CIP, EC_50_s of these drugs were quantified in the presence or absence of aTc. The ↓PSS-Dd2^A211V^ and ↓PSS-Dd2^G223R^ parasites were washed three times to remove residual aTc and then split into two cultures, one with aTc to maintain normal levels of *Pf*PSS and one without aTc to induce knockdown of *Pf*PSS levels. The cultures were allowed to grow for 48 hrs and then were seeded at 1% parasitemia and 1.5% hematocrit in a 96-well flat bottom well plate. The ↓PSS-Dd2^A211V^ parasites were introduced to a 1.5-fold dilution series of PA92 (a gift from Dr. Akhil Vaidya) (23.4 nM to 3.04 μM) and the ↓PSS-Dd2^G223R^ parasites were treated with a 3-fold dilution series of CIP (16.9 pM to 1 μM). The parasites were cultured for 72 hrs at 37 °C in a modular incubator chamber and growth was monitored by flow cytometry as described above.

For all the EC_50_ assays, parasitemia values from samples incubated with 0.1% DMSO (vehicle control) were used to calculate the relative parasite survival percentage. Percent survival data were collected from two independent biological replicates, each in quadruplicate, and were fit to a four-parameter sigmoidal dose-response curve in Prism V10.6 (GraphPad Software) to calculate the EC_50_ value. Statistical significance was determined using a two-tailed unpaired t-test with Welch’s correction.

## Data availability

WGS data for ↓PSS-Dd2^G223R^ parasites are available at the NCBI Sequence Read Archive (SRA) under submission PRJNA1426026.

## Acknowledgements

We are grateful to Akhil B. Vaidya (Drexel University) for his generous gifts of the Dd2 and Dd2^A211V^ parasite lines, and the PA21A092 and cipargamin inhibitors. We appreciate Anne Jedlicka and Amanda Dziedzic of the Johns Hopkins Bloomberg School of Public Health Genomic Analysis and Sequencing Core for their assistance in preparing and running samples for whole genome sequencing. We thank the Johns Hopkins Malaria Research Institute (JHMRI) Parasite Core for supplying O^+^ human red blood cells for parasite culture. This work was supported by the Network Science Initiative of the U.S. Army Medical Research and Development Command (USAMRDC) award W81XWH-20-C-0031 (STP), the National Institutes of Health (NIH) grant R01 AI125534 (STP), the Johns Hopkins Malaria Research Institute, and Bloomberg Philanthropies. AKM was supported by JHMRI predoctoral fellowship, MLS by NIH training grant T32AI007417, RE by JHMRI postdoctoral fellowship and Samuel Jordan Graham postdoctoral fellowship, and KR was supported by the Johns Hopkins Discovery Award. The funding agencies did not have any role in the design of the study, the collection or analysis of data, the decision to publish the findings, or the preparation of the manuscript.

