## Supplemental Information for "Reduced expression of an essential blood-stage *Plasmodium* phosphatidylserine synthase does not modulate parasite resistance to *Pf*ATP4 inhibitors"

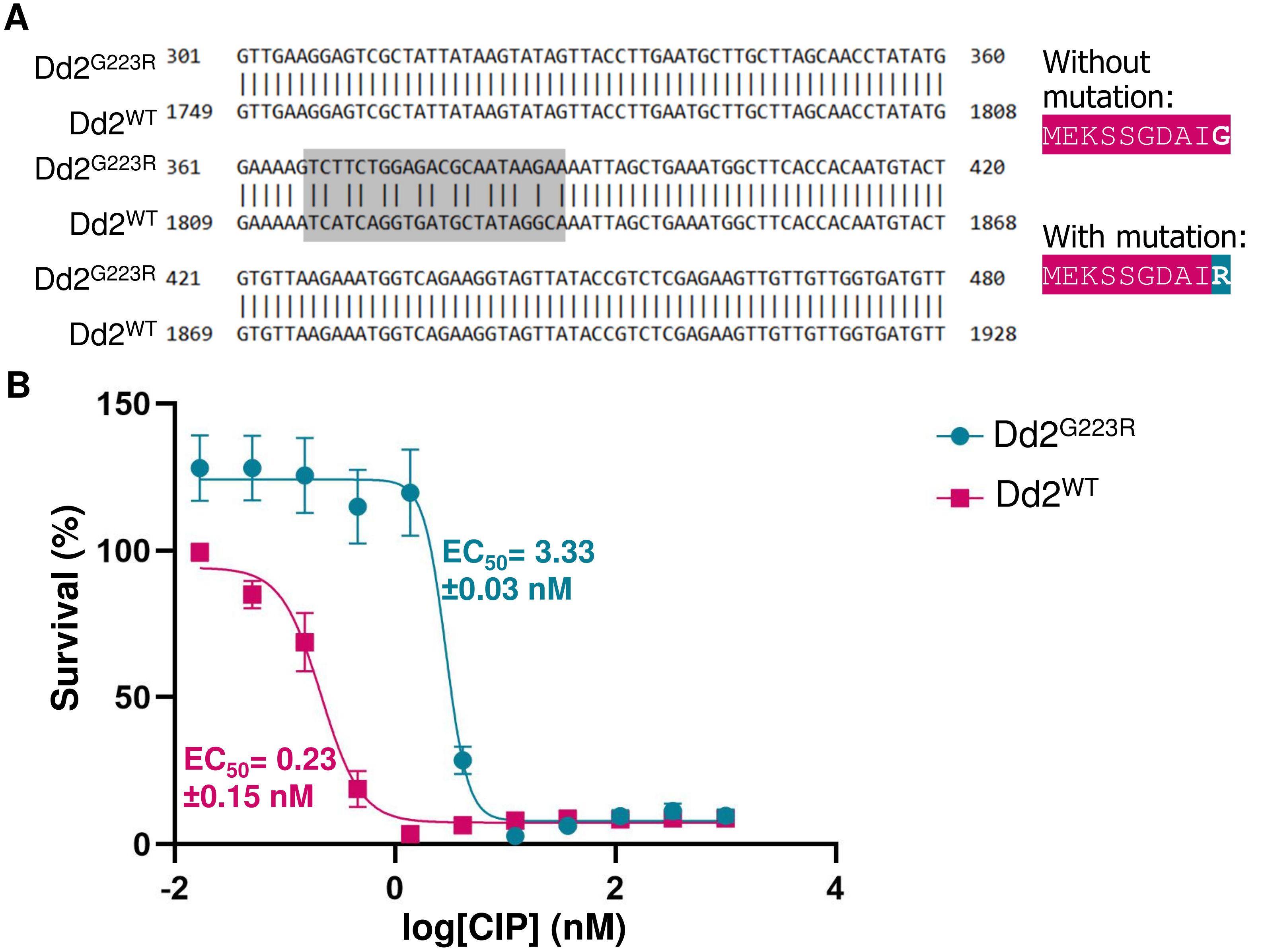

**Supplemental Figure 1. Cas9-mediated genetic editing of the *PfATP4* locus with a G223R mutation confers CIP resistance to Dd2 parasites.** A) Alignment of Sanger sequencing results from a Dd2$^{G223R}$ culture against the reference Dd2$^{WT}$ sequence. The gray boxed section of the alignment emphasizes the mutations incorporated into the *PfATP4* locus. This region includes shield mutations to prevent Cas9 cleavage of the mutated sequence. Except for the GCA → GAA single nucleotide polymorphisms (SNP) that results in the Gly223Arg amino acid replacement (highlighted in blue), the remaining SNPs do not result in a change to the coded amino acid (highlighted in pink). B) Relative survival curves for Dd2$^{WT}$ and Dd2$^{G223R}$ mutant parasite lines treated with a dilution series of CIP over the course of three days. Parasitemia at each drug dilution was normalized to the DMSO vehicle control as part of the % Survival calculations. The logarithmic nonlinear fit for each parasite line was determined in GraphPad Prism as log(inhibitor) vs. response with a variable slope (Dd2$^{G223R}$ $R^2$=0.8540; Dd2$^{WT}$ $R^2$=0.9033). Mean $EC_{50}$ values ± the standard deviation are shown. Error bars represent the standard error of the mean for two biological replicates, each conducted in quadruplicate.

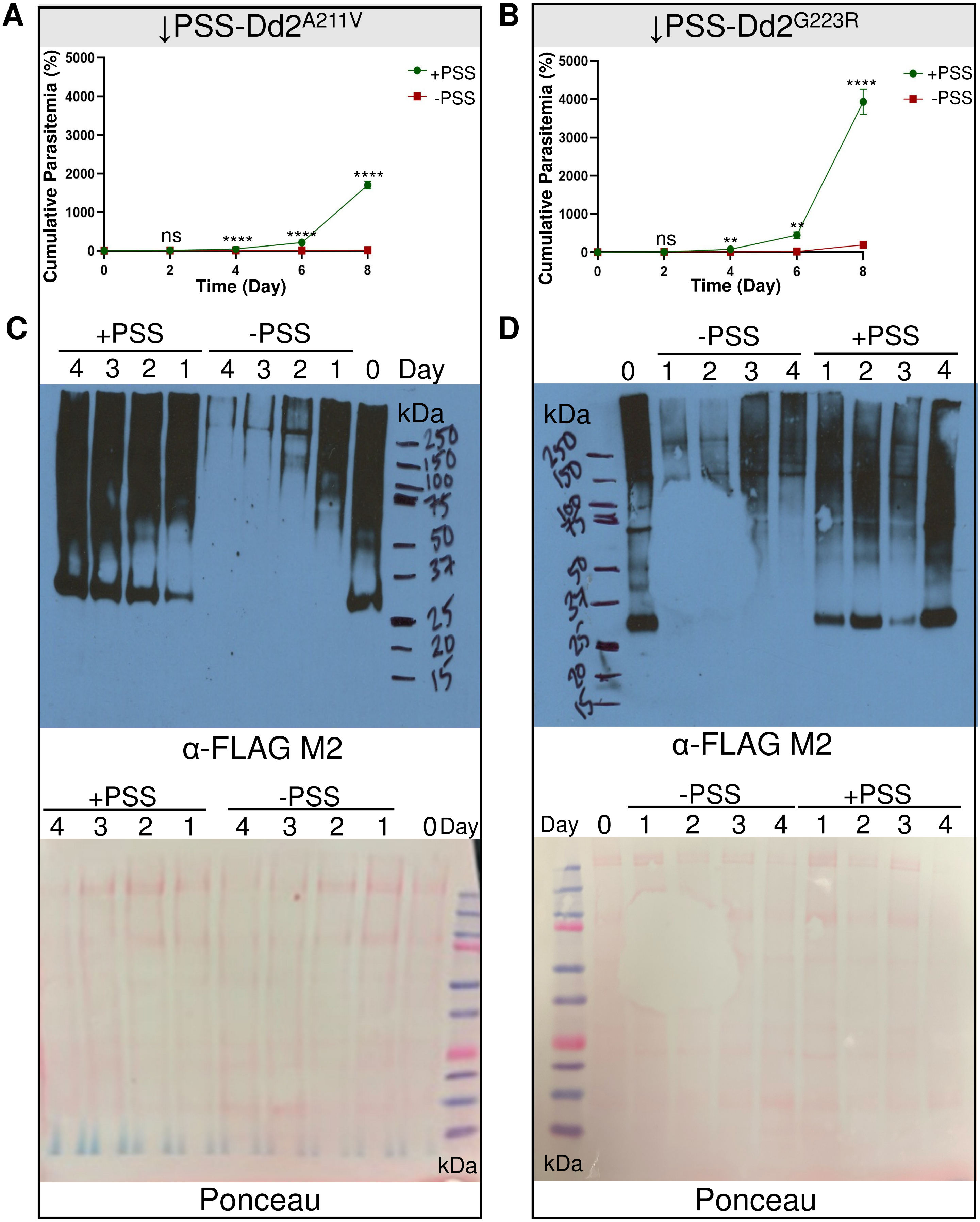

**Supplemental Figure 2. Reduction in *Pf*PSS protein levels leads to undetectable parasite growth.** Cumulative parasitemia graph depicting parasite growth for A) ↓PSS-Dd2$^{A211V}$ or B) ↓PSS-Dd2$^{G223R}$ cultured with aTc (+PSS) or without aTc (-PSS) over the course of eight days presented in Figure 3B and 3C, respectively. Parasites cultured with aTc have normal *Pf*PSS levels and grow with consistent culture cut-backs every two days. Parasites cultured without aTc have reduced *Pf*PSS levels and, by day four without aTc, begin showing statistically significant growth defects that result in undetectable parasite growth compared to those parasites grown in the presence of aTc. Error bars represent the standard error of the mean of two biological replicates, each conducted in quadruplicate. Statistical significance was determined using a two-way ANOVA with Šídák's multiple comparisons test. ns: not significant ($p>0.05$), ** significant ($p<0.009$), **** significant ($p<0.0001$). C) Uncropped versions of the Western blots for the ↓PSS-Dd2$^{A211V}$ or D) ↓PSS-Dd2$^{G223R}$ lines cultured either with aTc (+PSS) or without aTc (-PSS) as presented in Figure 3D and 3E, respectively. Bands for PSS, approximating 30 kDa, are visible when aTc is present (+PSS), but disappear by Day 1 following the removal of aTc from the parasite culture (-PSS). The FLAG-tagged PSS bands were identified following incubation with an α-FLAG primary antibody and the loading control was determined via whole-protein Ponceau staining. Large band sizes likely correspond to aggregated complexes of tagged protein.

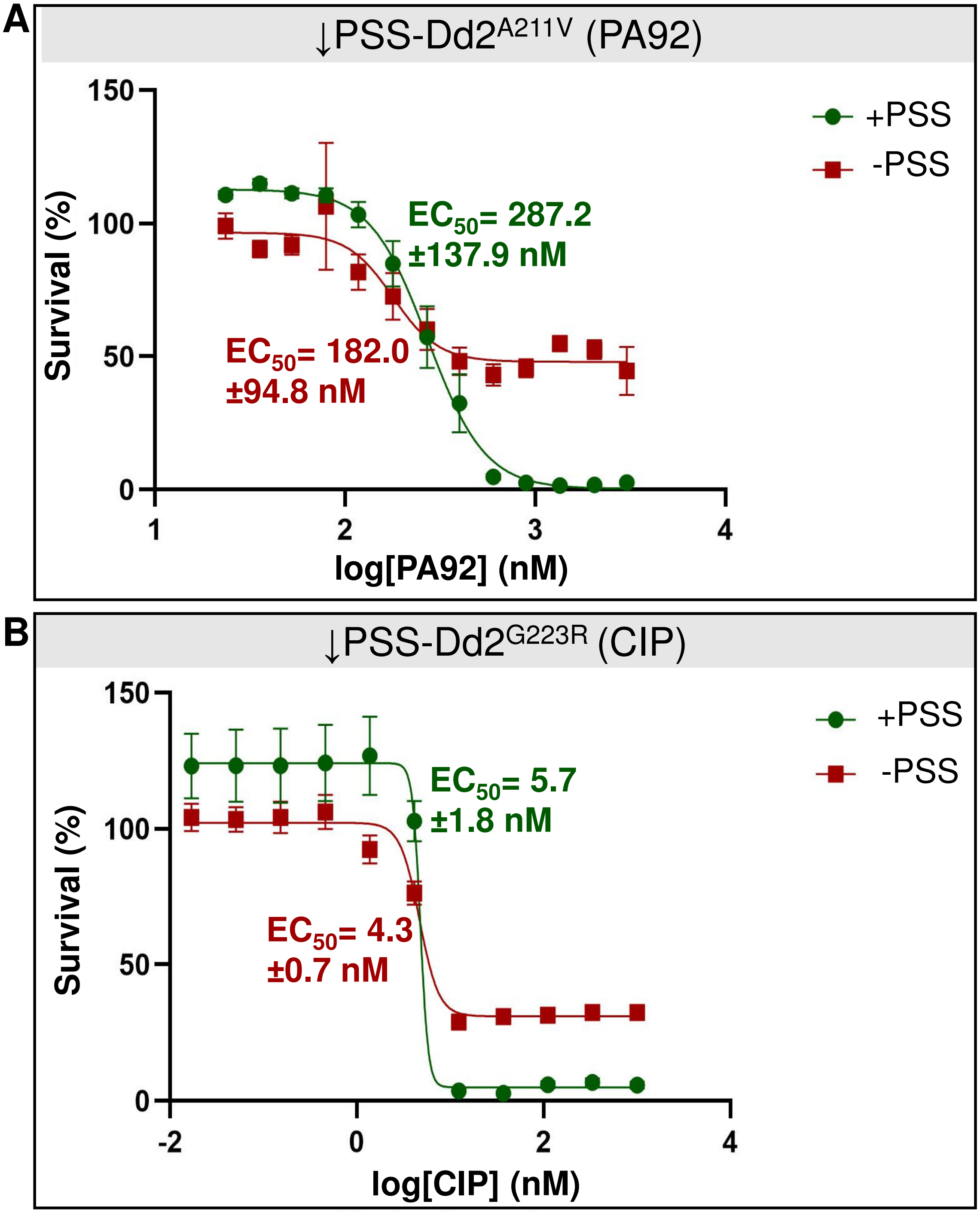

**Supplemental Figure 3. Reduced PSS levels do not alter response in the ↓PSS-Dd2$^{A211V}$ or ↓PSS-Dd2$^{G223R}$ parasite to PA92 or CIP, respectively.** Relative survival curves of the $EC_{50}$ assays for A) ↓PSS-Dd2$^{A211V}$ cultured either with normal or reduced PSS levels in the presence of a PA92 dilution series, or B) ↓PSS-Dd2$^{G223R}$ cultured either with normal or reduced PSS levels in the presence of a CIP dilution series. Parasitemia at each drug dilution was normalized to the DMSO vehicle control as part of the % Survival calculations. The logarithmic nonlinear fit for each parasite line was determined in GraphPad Prism as log(inhibitor) vs. response with a variable slope (↓PSS-Dd2$^{A211V}$ +PSS $R^2$=0.9199, -PSS $R^2$=0.4561; ↓PSS-Dd2$^{G223R}$ +PSS $R^2$=0.8437, -PSS $R^2$=0.9012). Mean $EC_{50}$ values ± the standard deviation are shown. Error bars represent the standard error of the mean of two biological replicates, each conducted in quadruplicate.

**Supplemental Table 1.** Primers and associated sequences used in this study. Restriction enzyme sites are underlined in the relevant primer sequences.

| Primer Name | Primer Description | Primer Sequence (5' → 3') | Restriction Enzyme Site |
|---|---|---|---|
| Complementary oligonucleotides targeting *PfATP4* G223 position | | | |
| G223R.gRNAF | Forward oligonucleotide for gRNA targeting of *PfATP4* G223 position | ATTAAGTATATAATATTAAATCATCAGGTGATGCTATGTTTTAGAGCTAGAAAT | |
| G223R.gRNAR | Reverse oligonucleotide for gRNA targeting of *PfATP4* G223 position | ATTTCTAGCTCTAAAACATAGCATCACCTGATGATTTAATATTATATACTTAAT | |
| Primers to amplify HAs for *Pf*PSS knockdown construct | | | |
| PSSb.HA1b.Bsa.F | Forward primer for *PfPSS* HA1 amplification | GGTGGTCTCGGATCCTTGATGTGTTGAGCTGTAACC | *Bsa*I-HF |
| PSSb.HA1b.Bsa.R | Reverse primer for *PfPSS* HA1 amplification | GGTGGTCTCAAATCCTATGGCAAATGTTCTAGCTATAACTATTAAATTTAC | *Bsa*I-HF |
| PSSb.HA2b.Asc.F | Forward primer for *PfPSS* HA2 amplification | GGTGGCGCGCCGGAGTTCCTTTTAGCCATTCGATGG | *Asc*I |
| PSSb.HA2b.Xho.R | Reverse primer for *PfPSS* HA2 amplification | GGTGGATCCCTCGAGGATATCAGAAGGAGTAGTATTGATGACAAAGTTG | *Xho*I |
| Oligonucleotide for *PfPSS* gRNA for gene knockdown | | | |
| PSSb.TD.gRNA | | cttacCAGAAATAAAACCTA | |

| | | |
|---|---|---|
| Primers to confirm integration of gene knockdown machinery into native locus | | |
| PSSb.5F | Forward primer for Δ5' and P PCRs | CATCCTATTCACACCGTGTGTTAGATATTCC |
| NewApt5R | Reverse primer for Δ5' PCRs | CTCGCTATCAAGGAATCGAGTCC |
| pTD.3F | Forward primer for Δ3' PCRs | CCAATGGCCCCTTTCCGGG |
| PSSb.3R | Reverse primer for Δ3' and P PCRs | GAAATGAAATGAAAAGAAAATTATTTAATCCCTTCATTG |
| Detection of mycoplasma contamination | | |
| Myco481.F | | CGAACTGAGATCGGCTTTTTGAG |
| Myco1250.R | | CCGCGGTAATACATAGGTCGC |
